# A D-alanine aminotransferase S_180_F substitution confers resistance to β-chloro-D-alanine in *Staphylococcus aureus* via antibiotic inactivation

**DOI:** 10.1101/2025.08.17.668425

**Authors:** Rakesh Roy, Yahani P. Jayasinghe, Sasmita Panda, Merve S. Zeden, Vinai C. Thomas, Donald R. Ronning, James P. O’Gara

**Affiliations:** Microbiology, School of Biological and Chemical Sciences, University of Galway, Ireland; Department of Pharmaceutical Sciences, University of Nebraska Medical Center, Omaha, Nebraska, USA; Fred & Pamela Buffett Cancer Center, University of Nebraska Medical Center Omaha Nebraska 68198 USA.; UNMC Center for Drug Design and Innovation, University of Nebraska Medical Center Omaha Nebraska 68198 USA.; Department of Pathology and Microbiology, University of Nebraska Medical Center, Omaha, Nebraska, USA

**Keywords:** alanine metabolism, alanine auxotrophy, BCDA resistance, D-alanine aminotransferase, crystal structure, alanine racemase, MRSA

## Abstract

Alanine transport and metabolism impact MRSA pathophysiology by dictating the availability of D-alanine for cell wall synthesis, the target of β-lactam antibiotics. Furthermore *cycA*-dependent alanine transport controls MRSA β-lactam susceptibility in chemically defined medium (CDM) in a glucose-dependent manner. Here we report that *S. aureus* was auxotrophic for L-alanine in CDM, and that this growth defect was rescued by glucose (or compensatory mutations), but only when the alanine racemase (*alr1*) and D-alanine aminotransferase (*dat*) genes were functional. No role was observed for the alanine dehydrogenase 1 (*ald1*) and *ald2* genes. As previously reported, *alr1* and, to a lesser extent, *cycA* mutations increased susceptibility to D-cycloserine (DCS). In contrast, only *alr1* mutation increased susceptibility to β-chloro-D-alanine (BCDA), suggesting distinct targets for these alanine analogue antibiotics, which act synergistically against MRSA. Genome sequencing of a BCDA-resistant mutant identified a C_539_T mutation in *dat*, predicted to result in a S_180_F substitution. Expression of the *dat*_C539T_ operon in wild-type increased BCDA resistance. *alr1/dat::Em* and *alr1/dat*_C539T_ double mutants were auxotrophic for D-alanine, indicating that Dat-S_180_F transaminase activity is impaired, a conclusion supported by *in vitro* enzyme assays. Structural modeling revealed an active-site loop shift in Dat-S_180_F that altered PLP co-factor binding. Molecular docking showed that the S_180_F substitution promotes BCDA-PLP adduct dissociation by releasing inactivated BCDA, thereby conferring resistance. These data reveal essential roles for Alr1 and Dat during growth under nutrient-limiting conditions and the potential of combination therapy separately targeting both enzymes with DCS and BCDA to extend the treatment options for MRSA infections.

## Introduction

According to the Centers for Disease Control and Prevention, *Staphylococcus aureus* is a leading cause of healthcare-associated infections in the United States where in 2017, it caused more than 119,000 bloodstream infections and nearly 20,000 deaths (1). Methicillin-resistant *S. aureus* (MRSA) has been designated a high-priority pathogen by the World Health Organization in both its 2017 and 2024 Bacterial Priority Pathogens Lists, highlighting the ongoing and urgent need for novel, effective antimicrobial therapies targeting MRSA infections (2). The emergence of antibiotic resistance in *S. aureus* poses a critical threat to global public health, contributing significantly to the burden of AMR, with over 100,000 deaths attributable to MRSA in 2019 alone (3, 4). *S. aureus* is well-known for its ability to acquire resistance to almost all currently available antibiotics. Among these, its resistance to β-lactam antibiotics, a cornerstone in the bacterial infection treatment, is particularly alarming. Resistance to methicillin and other β-lactams in *S. aureus* is conferred by the acquisition of the *mecA* gene (5). This gene encodes penicillin-binding protein 2a (PBP2a), a transpeptidase enzyme that exhibits reduced affinity for β-lactam antibiotics, allowing bacterial cell wall crosslinking in the presence of these drugs. However, β-lactam resistance in *S. aureus* is not solely driven by *mecA*, and additional genetic and metabolic factors also play a role in modulating this resistance phenotype (6–11). Understanding these complex resistance mechanisms is an important part of efforts to identify and target enzymes and pathways that may resensitise MRSA to β-lactam antibiotics.

Bacterial cell wall biosynthesis is an important target of many antibiotics, and altering the physiology of the cell wall by targeting pathways associated with it can re-sensitize MRSA to β-lactam antibiotics (12, 13). An important constituent of the cell wall is D-alanine, which is incorporated into the peptidoglycan stem peptide. Alanine transport via CycA and the pathway leading to the incorporation of the D-ala-D-ala residues into the peptidoglycan stem peptide plays an important role in the susceptibility of MRSA to β-lactam antibiotics (14, 15). Our previous work showed that mutation of *cycA* or MRSA exposure to the alanine/serine analogue D-cycloserine (DCS), which blocks the activity of the D-alanine racemase (Alr) and D-alanine ligase (Ddl) enzymes, significantly potentiated oxacillin (OX) *in vitro* and in a mouse model of bacteremia (14). However, these phenotypes were only evident in complex media (Mueller Hinton Broth), or chemically defined media supplemented with glucose (CDMG). In CDM lacking glucose (CDM), alternative carbon sources such as amino acids become more important for growth (16), and the impact of the *cycA* mutation on DCS or OX susceptibility was significantly ameliorated (14), suggesting that an alternative alanine transporter(s) or altered alanine metabolism may compensate for CycA under these growth conditions. In particular, endogenous L- and D-alanine biosynthetic pathways may play a more significant role in the absence of CycA-mediated alanine transport in CDM. L-alanine can be synthesised from pyruvate by alanine dehydrogenase 1 and 2 (Ald1 and Ald2), while D-alanine can be synthesized from pyruvate and D-glutamate by D-alanine aminotransferase (Dat) (Fig. 1). A *S. epidermidis* triple mutant lacking *alr1*, *alr2* and *dat* was previously shown to be auxotrophic for D-alanine (17). Notably exposure of MRSA to sub-inhibitory OX increased transcription of both *ald1* and *dat* (18), which appears to be consistent with an increased requirement for alanine under β-lactam stress. Increased Ald1 or Dat activity may increase L-alanine and D-alanine biosynthesis, respectively, irrespective of reduced alanine transport in CycA or other putative alanine transporter mutants. The role of Alr2 which shares 29% homology with Alr1 remains unclear (19). Synergy between DCS and another alanine analogue antibiotic, β-chloro-D-alanine (BCDA) and β-lactams has previously been reported (20), but potential synergy between DCS and BCDA has not been investigated. BCDA has been reported to inhibit Alr1 activity in a number of Gram negative organisms (21–23), but not *Mycobacterium tuberculosis* (24) and the mechanism of action of BCDA in *S. aureus* remains to be determined.

**Fig. 1.**
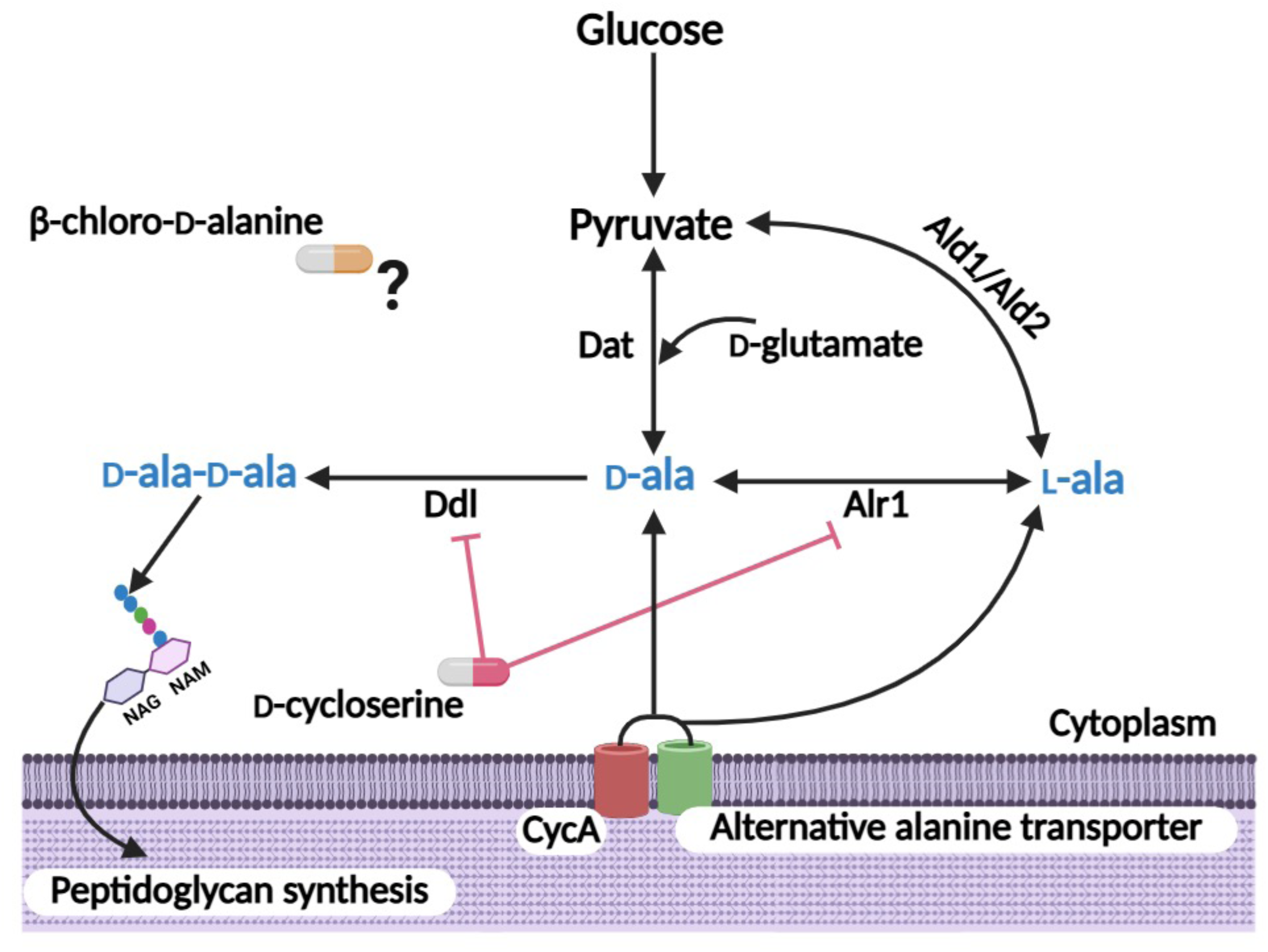
Schematic depicting the major enzymes involved in production of the D-ala-D-ala dipeptide required for peptidoglycan (PG) biosynthesis in *S. aureus*. Our previous data have implicated CycA or an alternative transporter(s) in alanine uptake. Alanine racemase Alr1 converts L-alanine (L-ala) to D-alanine (D-ala), and D-ala-D-ala is synthesized by D-alanine ligase (Ddl). Within the cell L-alanine can be synthesized from pyruvate by alanine dehydrogenase 1 and 2 (Ald1 and Ald2), while D-alanine can be synthesized from pyruvate and D-glutamate by D-alanine aminotransferase (Dat). D-cycloserine blocks the activity of the Alr1 and Ddl enzymes. The antimicrobial mode of action of β-chloro-D-alanine in *S. aureus* remains largely unexplored. Created with Biorender.com.

In this study, we examined the impact of mutations in the D-ala-D-ala and endogenous alanine biosynthetic pathways on MRSA susceptibility to OX, DCS and BCDA. An *in vitro* evolution experiment identified Dat as a target for BCDA and a C_539_T single nucleotide substitution resulting in a predicted S_180_F amino acid substitution was implicated in increased BCDA resistance. An *alr1/dat*_C539T_ mutant was constructed and the enzymatic activity of purified recombinant Dat-S_180_F was measured. The impact of the S_180_F substitution on Dat activity was modelled after determining the X-ray crystal structure of Dat and its pyridoxal 5’-phosphate (PLP) co-factor. BCDA is expected to undergo β-elimination upon reacting with PLP to form a covalent complex with Dat residue Lys146 and irreversibly inactivating the enzyme (25). However, our data suggests that Dat-S_180_F may bind and react with BCDA but avoid covalent modification of the essential active site lysine residue. Combinations of β-lactams, BCDA (targeting Dat) and/or DCS (targeting Alr1 and Ddl) may have therapeutic potential to improve the treatment of MRSA infections.

## Results

### Reversal of MRSA L-alanine auxotrophy by glucose or compensatory mutations is dependent on alanine racemase (Alr1) and D-alanine aminotransferase (Dat)

Alanine transport and the D-ala-D-ala pathway control MRSA susceptibility to β-lactams and early cell wall inhibitors such as DCS in a culture medium dependent manner (14). Building on our previous observation that growth in chemically defined medium (CDM) compared to CDM with glucose (CDMG) or Mueller Hinton broth (MHB) increased β-lactam and DCS resistance in the *cycA* alanine transport mutant (14), we compared growth of wild-type JE2 and the alanine metabolism mutants *alr1*, *alr2*, *ald1*, *ald2, cycA* and *dat* in CDM (Fig. 2A, B) and CDMG (Fig. 2C, D). A *ddl* mutant is not available in the Nebraska Transposon Mutant Library (NTML). JE2 is auxotrophic for L-alanine in CDM (Fig. 2B) but not CDMG (Fig. 2D) consistent with an important role for carbon flux from central metabolism to the D-ala-D-ala pathway in the absence of exogenous alanine. However, it is notable that, apart from *alr1* and *dat*, weak growth was detected for JE2, *cycA*, *alr2*, *ald1* and *ald2* after a 16-18 h lag in CDM lacking L-alanine (Fig. 2B). Faster growth of these strains after subculture in fresh CDM lacking L-alanine (data not shown) suggested the acquisition of compensatory mutations that perhaps enhance the flux of alternative amino acids into central metabolism in the absence of L-alanine. The analysis of JE2 revertants to L-alanine prototrophy will be the subject of a separate study.

**Fig. 2.**
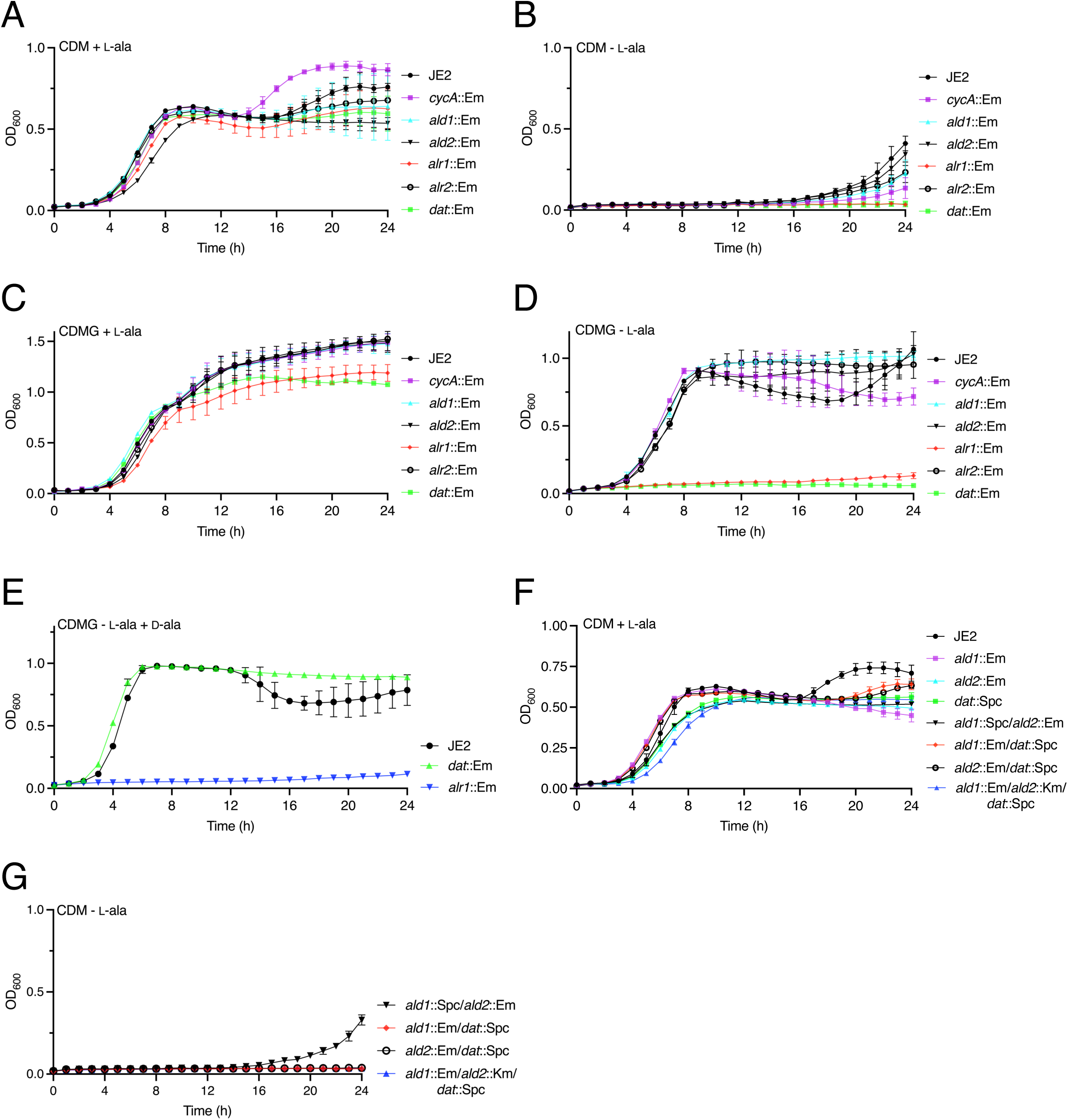
*S. aureus* is auxotrophic for L-alanine in CDM but not CDMG. **A - D.** Comparison of JE2, *cycA* (*cycA*::Em, NE810), *ald1* (*ald1*::Em, NE1136), *ald2* (*ald2*::Em, NE198), *alr1* (*alr1*::Em, NE1713, *alr2* (*alr2*::Em, NE799) and *dat* (*dat*::Em, NE1305) growth in CDM + L-alanine (A), CDM - L-alanine (B), CDMG + L-alanine (C) and CDMG - L-alanine (D). **E.** Comparison of JE2, *alr1* and *dat* growth in CDMG - L-alanine + D-alanine. **F.** Comparison of JE2, *ald1, ald2, dat, ald1*::Spc*/ald2*::Em*, ald1*::Em/*dat::*Spc*, ald2*::Em/*dat*::Spc and *ald1*::Em/*ald2*::Km/*dat::*Spc growth in CDM + L-alanine (G). **G.** Comparison of JE2, *ald1*::Spc*/ald2*::Em*, ald1*::Em/*dat::*Spc*, ald2*::Em/*dat*::Spc and *ald1*::Em/*ald2*::Km/*dat::*Spc growth in CDM - L-alanine. Where indicated L- and D-alanine were added at a final concentration of 5mM. The data presented are the average of at least 3 biological replicates and standard deviations are shown.

In CDMG, L-alanine was required for growth of the *alr1* (but not *alr2*) and *dat* mutants (Fig. 2C,D). Presumably Dat is required for the synthesis of D-alanine from pyruvate in CDMG lacking L-alanine (Fig. 1). However, it was less clear why Dat cannot compensate for *alr1* under these growth conditions. One possibility is that Alr1 is required for conversion of D-alanine produced by Dat to L-alanine in CDMG lacking L-alanine. Supporting this idea, the growth defect of *alr1* in CDMG lacking L-alanine (Fig. 2D) was not rescued by D-alanine alone (Fig. 2E).

Compared to JE2, mutations in *ald1* and *ald2* did not impact growth under the conditions tested (Fig. 2A-D) including in CDMG lacking L-alanine (Fig. 2D), suggesting that these enzymes are not required for L-alanine biosynthesis or the D-ala-D-ala pathway when growing on glucose. Conversely, we recently demonstrated that L-alanine is fluxed to pyruvate during growth in CDM (19), indicating that Ald1 and/or Ald2 are involved in the conversion of L-alanine to pyruvate when this amino acid is available as a carbon source. However, as noted above, *ald1* and *ald2* mutants grew normally in CDM (Fig. 2A). Furthermore, to rule out the possibility that Ald1 and Ald2 can compensate for each other, an *ald1*/*ald2* double mutant was constructed and also shown to grow normally in CDM (Fig 2F). To investigate if Dat can play a role in compensating for the Ald enzymes, *ald1*/*dat, ald2*/*dat* and *ald1*/*ald2*/*dat* mutants were constructed. All three of these mutants grew normally in CDM (Fig. 2F) and all were growth impaired in CDM - L-alanine (Fig. 2G). Interestingly as observed for *ald1* and *ald2* single mutants (Fig. 2B), the *ald1*/*ald2* double mutant did start to grow after a 16 h lag in CDM - L-alanine (Fig. 2G) but all combination mutants lacking *dat* showed no growth even after 24 h (Fig. 2G). While the functions of Ald1 and Ald2 in the D-ala pathway remain unclear, these data do reveal that reversal of MRSA L-alanine auxotrophy in CDM by glucose or compensatory mutations is dependent on Dat-mediated conversion of pyruvate to D-alanine, and the subsequent conversion of D-alanine to L-alanine by Alr1.

### Mutation of *alr1* but not *cycA* increases β-chloro-D-alanine (BCDA) susceptibility in CDM

Consistent with our previous data (14), the *alr1* mutant was dramatically more susceptible to OX and DCS in CDM (Table 1). The *cycA* mutation also increased susceptibility to OX and DCS (Table 1), albeit to a lesser extent to that previously observed in CDM supplemented with glucose (14). Interestingly, while the *alr1* mutation significantly increased susceptibility to another alanine analogue antibiotic BCDA, the *cycA* mutation had no effect (Table 1). The susceptibility of the *ald1* and *ald2* mutants to OX, BCDA and DCS was not significantly different to wild-type (Table 1). Mutation of *dat* marginally reduced the BCDA MIC from 300 to 200 μg/ml and had no effect on OX or DCS susceptibility (Table 1). The *alr2* mutant had wild-type levels of OX, DCS and BCDA susceptibility (Table 1). Taken together these data support important roles for Alr1 and Dat in the susceptibility of MRSA to DCS and BCDA. The differential effect of the *cycA* mutation on DCS and BCDA susceptibility suggests that these antibiotics have different targets or mechanisms of action in *S. aureus*. Furthermore, disk diffusion and checkerboard titration assays (ΣFIC = 0.17) revealed synergy between DCS and BCDA when used in combination against JE2 (Fig. S1A,B).

**Table 1.** Antibacterial activity (minimum inhibitory concentrations, MICs; μg/ml) of D-cycloserine (DCS), oxacillin (OX) and β-chloro-D-alanine (BCDA) against wild-type MRSA strain JE2 and alanine metabolism mutants grown in chemically defined media lacking glucose (CDM).

| Strain | DCS | OX | BCDA |
| --- | --- | --- | --- |
| JE2 | 32 | 32 | 300 |
| NE810 (*cycA*) | 4 | 8 | 300 |
| NE1713 (*alr1*) | <0.5 | <0.5 | <50 |
| NE799 (*alr2*) | 32 | 32 | 300 |
| NE1136 (*ald1*) | 32 | 32 | 300 |
| NE198 (*ald2*) | 32 | 16 | 300 |
| NE1305 (*dat*) | 32 | 32 | 200 |
| NE1898 (*pepV*) | 32 | 32 | 300 |
| BCDA1 (*dat*<sub>C539T</sub>) | 32 | 32 | 800 |
| BCDA1 *dat*::Em | 32 | 32 | 300 |
| JE2 *ppepV-dat*<sub>C539T</sub> | 32 | 32 | 600-700 |

### A S_180_F substitution in D-alanine aminotransferase (Dat) confers resistance to β-chloro-D-alanine (BCDA) in *S. aureus*

To investigate the mechanism of action of BCDA, a single-step, high selective pressure experiment was used to isolate a stable BCDA resistant mutant of JE2. This mutant, designated BCDA1, was isolated from a 96 well plate in which JE2 was grown in CDM lacking glucose supplemented with an inhibitory concentration (500 μg/ml) of BCDA. Whole-genome sequencing identified a single nucleotide mutation, C-to-T, at position 539 of the *dat* gene (SAUSA300_1696) predicted to result in a S_180_F amino acid substitution (Fig. 3A). No other genetic changes were identified in BCDA1 and the *dat*_C539T_ mutation was confirmed by PCR amplification and Sanger sequencing.

**Fig. 3.**
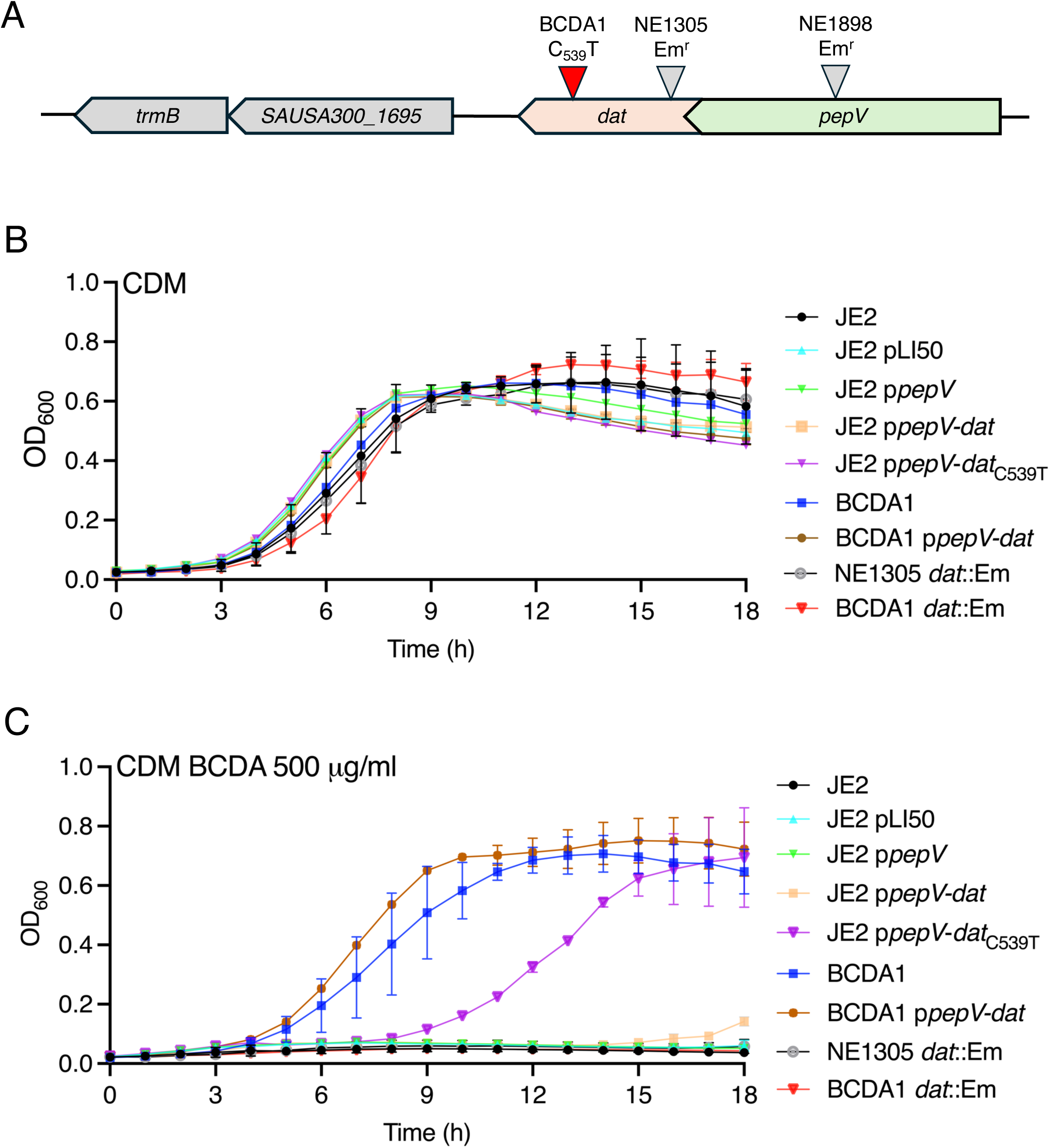
The *dat*_C539T_ mutation increases BCDA resistance in *S. aureus*. **A.** Genomic organisation of the *dat* locus in *S. aureus.* The 849 bp *dat* gene (SAUSA300_1696) is in a 2-gene operon with the dipeptidase gene *pepV*. A C_539_T mutation predicted to result in a S_180_F amino acid substitution was identified in the BCDA resistant strain BCDA1. The location of the Tn insertion in the NTML *dat* mutant NE1305 is also shown. **B and C.** Comparison of JE2, JE2 pLI50, JE2 p*pepV*, JE2 p*pepV*-*dat*, JE2 p*pepV*-*dat*_C539T_, BCDA1, BCDA1 p*pepV*-*dat*, NE1305 (*dat*::Em) and BCDA1 *dat*::Em growth in CDM (A) and CDM supplemented with BCDA 500 μg/ml (B). The data presented are the average of at least 3 biological replicates and standard deviations are shown.

The BCDA MIC for BCDA1 increased to 800 μg/ml compared to 300 μg/ml for wild-type JE2 and 200 μg/ml for NE1305 (*dat*::Em) (Table 1). In liquid CDM cultures, growth of BCDA1 was the same as JE2 and NE1305 (Fig. 3B), whereas in CDM supplemented with 500 μg/ml BCDA, only the resistant mutant was able to grow (Fig. 3C). Phage 80α-mediated transduction of the *dat*::Em allele from NE1305 into BCDA1 restored BCDA susceptibility (Fig. 3C) and reduced the BCDA MIC to wild-type levels (300 μg/ml) (Table 1). Multicopy expression of the 2-gene *pepV*-*dat*_C539T_ operon from BCDA1 increased BCDA resistance in CDM liquid cultures, albeit after a longer lag period (Fig. 3C) and increased the BCDA MIC to 600-700 μg/ml (Table 1). In contrast multicopy expression of the wild-type *pepV*-*dat* operon in JE2 had no significant effect on growth in CDM supplemented with BCDA (Fig. 3C). Multicopy expression of the *pepV* dipeptidase gene alone in JE2 for control purposes also had no effect on BCDA resistance (Fig. 3C). Collectively these data support the conclusion that the *dat*_C539T_ allele in BCDA1 is the critical determinant driving increased resistance to BCDA.

### BCDA1 is auxotrophic for D-alanine in the absence of *alr1*

Next the impact of the *dat*_C539T_ mutation in BCDA1 on the D-ala-D-ala pathway and alanine auxotrophy was investigated. Phage 80α was used to transduce the *alr1*::Em allele from NE1713 into BCDA1. Consistent with recently published data (17, 19), we hypothesized that in the absence of alanine racemase, synthesis of D-alanine will be entirely dependent on Dat activity (Fig. 1). The *alr1*/*dat*_C539T_ double mutant was unable to grow in TSB, unless the growth media was supplemented with D-alanine (Fig. 4A). Supplementation of TSB, which already contains enough L-alanine to support the growth of *alr1* (Fig. 4A), with additional L-alanine was unable to rescue growth of the *alr1*/*dat*_C539T_ mutant (Fig. 4A), further supporting the requirement for D-alanine. For control purposes an *alr1*/*dat* double mutant carrying transposon insertions in both genes was also constructed. To facilitate this, the *dat*::Em allele in NE1305 was first replaced with a Spc cassette and the *dat*::Spc mutant then transduced with the *alr1*::Em allele. Growth of the *alr1*::Em/*dat*::Spc double mutant in TSB was also dependent on D-alanine (Fig. 4A). Consistent with growth data indicating that Alr2 is not involved in the D-ala-D-ala pathway (Fig. 1), transduction of the *alr2*::Em allele from NE799 into BCDA1 had no effect on growth in TSB in the absence of D-alanine (Fig. 4A).The *alr1*/*dat*_C539T_ and *alr1*::Em/*dat*::Spc mutants were unable to grow in CDMG lacking L- and D-alanine (Fig. 4B) or supplemented with L-alanine (Fig. 4C) or D-alanine alone (Fig. 4D) but did grow when provided with both L- and D-alanine (Fig. 4E). Taken together, these data demonstrate that BCDA1 is auxotrophic for D-alanine in the absence of *alr1* and suggest that the Dat-S_180_F mutation negatively impacts D-alanine aminotransferase activity.

**Fig. 4.**
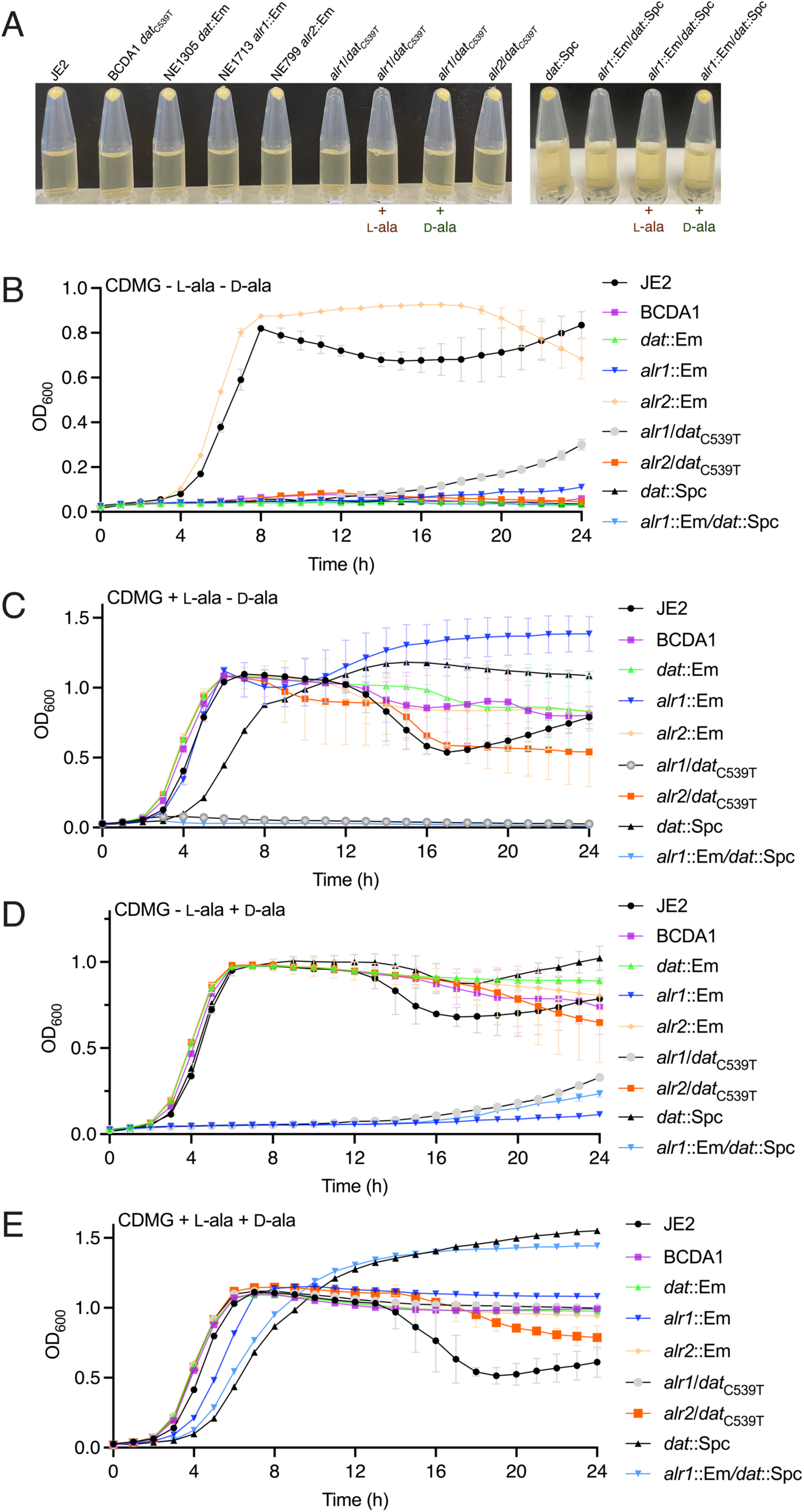
BCDA1 is auxotrophic for D-alanine in the absence of *alr1*. **A.** Growth of JE2, BCDA1 (*dat*_C539T_), NE1305 *dat*::Em, NE1713 *alr1*::Em, NE799 *alr2*::Em, *alr1*/*dat*_C539T_, *alr2*/*dat*_C539T_, *dat*::Spc and *alr1*::Em/*dat*::Spc for 18 h at 37°C in TSB supplemented as indicated with 5 mM L-alanine or D-alanine. Growth or no growth is indicated by the presence or absence of a cell pellet following microcentrifugation of 1 ml culture aliquots. This experiment was repeated 3 times, and the results of a representative experiment is shown. **B, C, D and E.** Comparison of JE2, BCDA1 (*dat*_C539T_), NE1305 *dat*::Em, NE1713 *alr1*::Em, NE799 *alr2*::Em, *alr1*/*dat*_C539T_, *alr2*/*dat*_C539T_, *dat*::Spc and *alr1*::Em/*dat*::Spc growth in CDMG with no L- or D-alanine (B) CDMG with 5mM L-alanine (C), CDMG with 5mM D-alanine (D) and CDMG with both L- and D-alanine (5mM) (E). The data presented are the average of at least 3 biological replicates and standard deviations are shown.

### The Dat-S_180_F variant has reduced activity and a lower IC_50_ for BCDA

The wild-type *dat* and *dat*_C539T_ genes were cloned into pET28b and HIS-tagged recombinant proteins were purified from *E. coli* via nickel affinity chromatography. To compare the activity of Dat and Dat-S_180_F, we quantified pyruvate produced by both proteins using a lactate dehydrogenase coupled assay. The activity of Dat-S_180_F is 43% of the wild-type level (Fig. 5A). Inhibition of Dat-S_180_F by BCDA displayed a lower IC_50_ (34.0 ± 4.7 µM; Fig. 5A) as compared to wild-type Dat (IC_50_ = 96.8 ± 12.0 µM). Similar to the *alr1*/*dat*::Spc mutant, the *alr1*/*dat*_C539T_ mutant was also auxotrophic for D-alanine (Fig. 4A-E) indicating that the reduced activity of the Dat-S_180_F variant is not sufficient to compensate for the absence of Alr1. Taken together, these data suggest that that the lower IC_50_ observed for the S_180_F variant is likely a consequence of the lower intrinsic enzymatic activity and that the inhibition levels observed at BCDA concentrations higher than 400 μM are more reflective of inhibition levels at the determined MICs.

**Fig. 5.**
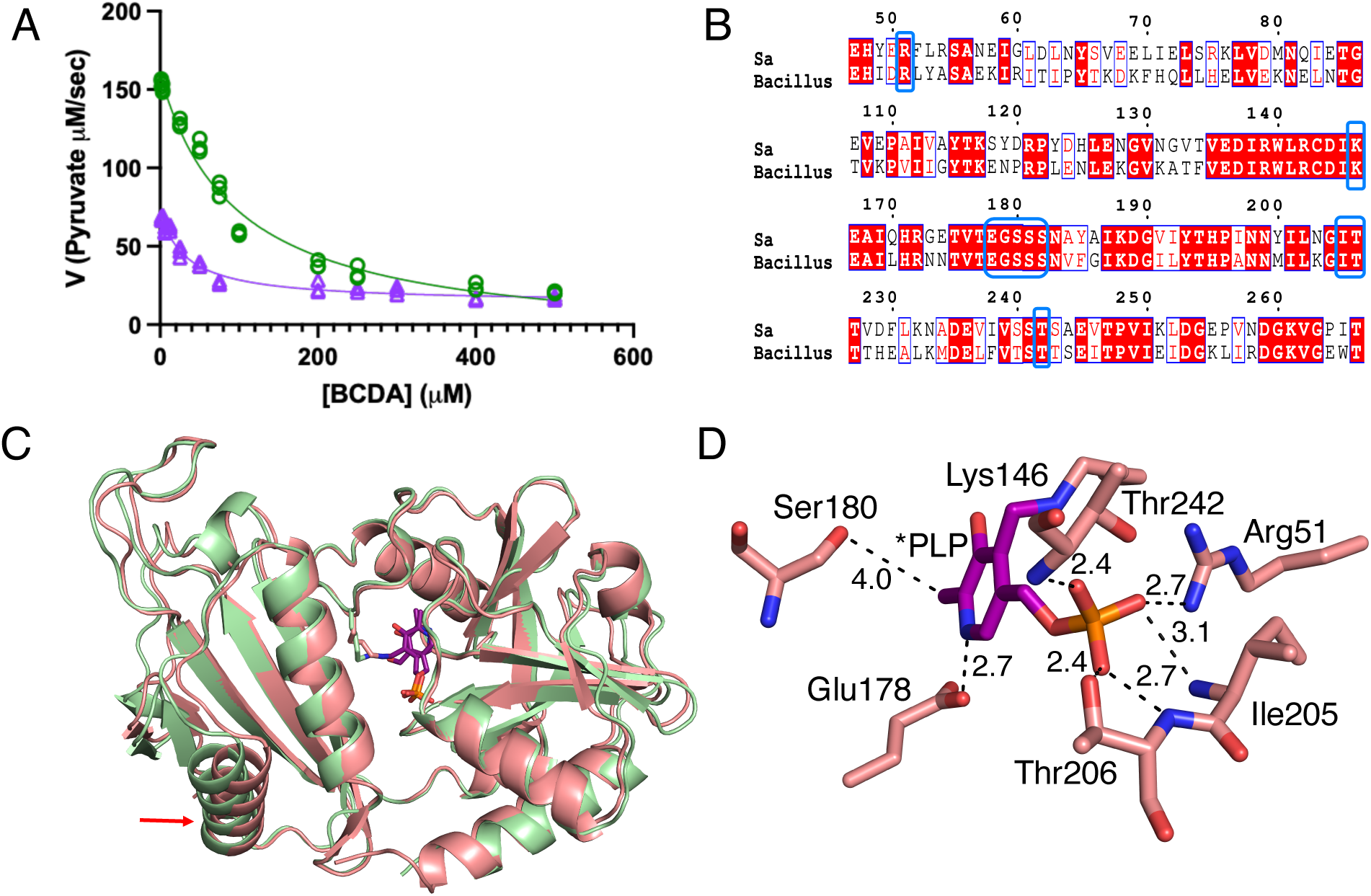
The S_180_F mutation reduces Dat enzymatic activity and increases the inhibitory activity of BCDA. **A.** Comparison of wild-type Dat (green) and Dat-S_180_F (violet) enzymatic production of pyruvate from D-alanine (which was coupled to that of lactate dehydrogenase) in increasing concentrations of BCDA (up to 500 μM). The enhanced inhibitory activity observed at BCDA concentrations below 400 μM is no longer evident at higher BCDA concentrations. **B.** Amino acid sequence alignment of the *S. aureus* (Sa) and *Bacillus* sp. Dat proteins. **C.** Superimposition of the *S. aureus* (salmon) and *Bacillus* sp. YM-1 (RCSB PDB: 1DAA) (green) Dat structures. The red arrow (bottom left) indicates the position of the structural differences. **D.** The interactions of PLP of aldimine with neighboring residues. The carbon atoms of PLP are in purple and the carbon atoms in the *S. aureus* Dat structure are in salmon color. The oxygen, nitrogen, and phosphate atoms are in red, blue, and orange, respectively.

### Structure of D-alanine aminotransferase in complex with its coenzyme pyridoxal phosphate (PLP)

To gain mechanistic insights into how the S_180_F mutation impacts Dat activity, the X-ray crystal structure of Dat was solved at 2.9 Å resolution in a P4_1_ 2_1_ 2 space group with one molecule in the asymmetric unit. In its native biologically relevant form, Dat is a dimer. The crystallographic 2-fold axis relates the asymmetric unit contents to the second Dat subunit of the biological dimer. The crystal structure shows that PLP forms the expected covalent aldimine with Lys146 as indicated by continuous electron density. The comparison of *S. aureus* Dat with the published structure of *Bacillus* sp. YM-1 Dat (RCSB PDB: 1DAA) reveals only minor structural differences even though the sequence identity between the two proteins is 52% (Fig. 5B). Only helix α1 and the preceding loop deviate between the two orthologs (Fig. 5C). The active site residues of both structures exhibit similar conformations and interactions with PLP.

The phosphate group of PLP interacts with the guanidium group nitrogen of Arg51, backbone nitrogen of Ile205, backbone nitrogen and side chain of Thr206 and backbone nitrogen of Thr242 by forming hydrogen bonds. The nitrogen of the pyridine forms an interaction with Glu178. The methyl group of PLP resides at 4.0 Å from the backbone carbonyl group of Ser180 forming a modest van der Waals interaction (Fig. 5D).

### Comparison of the Dat and Dat-S_180_F structures

As Dat possesses a shared active site with residues from both subunits contributing to PLP and substrate binding, this necessitates the use of a biological dimer for docking experiments. The biological dimer of Dat was made using crystallographic symmetry, and energy minimization was performed in Glide to have all the residues in the lowest energy conformation. The corresponding Dat-S_180_F model was made from the wild-type crystal structure with residue 180 in both subunits being mutated to Phe. This was again followed by the energy minimization in Glide. Compared to the 4 Å in the crystal structure, docking with covalent adduct produces a slight shift of the PLP that reduces the distance between the methyl group of PLP and the Ser180 to 3.5 Å. This is unlikely to reflect use of the slightly larger PLP adducts in the docking, because the same shift is observed when PLP is attached to catalytic residue Lys146, but illustrates the possible movement in the 180-182 loop that may impact PLP binding or affinity during various steps of the catalytic reaction. Notably, the side chains in this loop have already been identified as important residues for interactions with D-amino acid substrates or products in other homologous proteins (26). In contrast, comparison of the Dat and Dat-S_180_F dimer structures revealed no significant differences apart from the deviation of the loop harboring Phe180 (Fig. 6A).

**Fig. 6.**
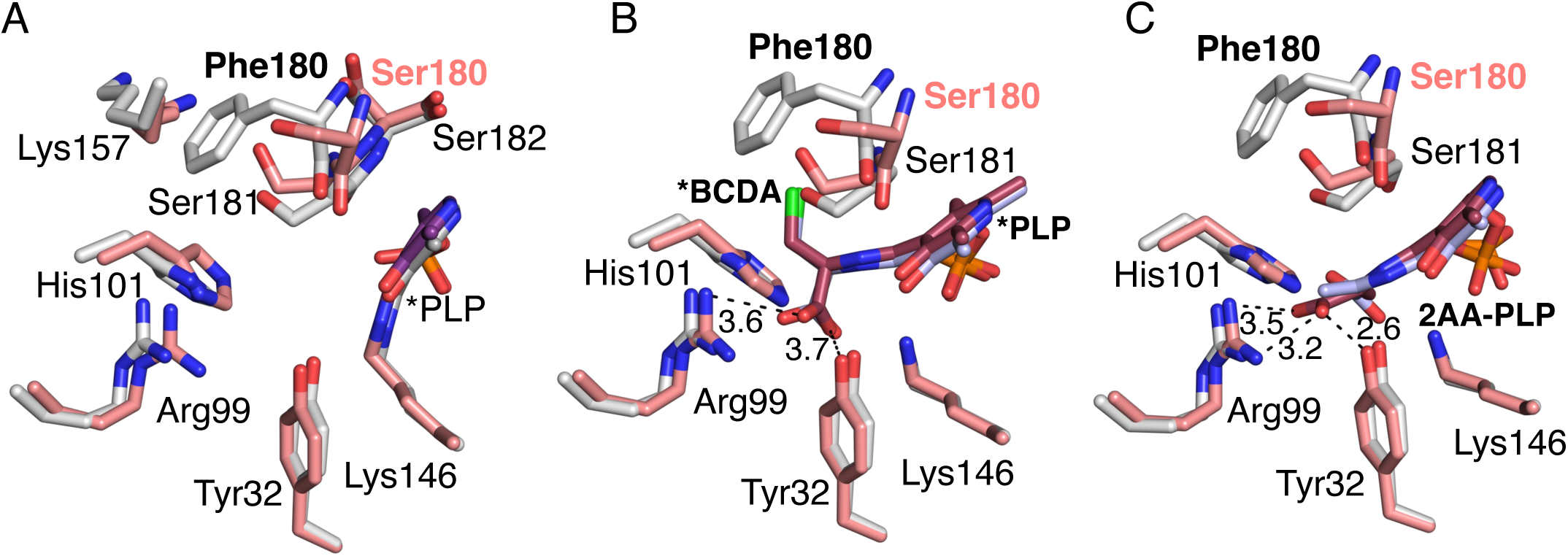
**A.** Conformational differences between amino acid residues in wild-type *S. aureus* Dat dimer structure (carbons in salmon) and Dat-S_180_F (carbons in gray). The nitrogen and oxygen atoms are in blue and respectively. The PLP is forming an internal aldimine with Lys146. **B.** The docking results of PLP-BCDA covalent adduct (external aldimine) to the wild-type Dat (carbons in salmon) and Dat-S_180_F (carbons in gray) variant indicating similar binding modes. **C.** Docking of the 2AA external aldimine. The carbons of the 2AA external aldimine produced by β-elimination of the PLP-BCDA adduct are in pink and grey in the wild-type Dat and Dat-S_180_F structures, respectively. The differing orientations of 2AA maybe be one way that the variant avoids covalent modification of Lys146.

This otherwise minor structural change has multiple effects that may impact PLP-BCDA affinity following formation of that adduct. The alteration in the loop harboring residue 180 increases the distance between the methyl group of PLP with the backbone of Phe180 versus Ser180, resulting in a shift from 3.5 to 4.5 Å. As the aromatic ring of PLP also forms 7f-7f interactions with the peptide bond between residues 181 and 182, this minor structural shift due to the S_180_F mutation could result in a larger than expected affinity change. Also contributing to a possible affinity decrease, the *B. subtilis* Dat structure (RSCB PDB:3DAA) in a complex with a PLP-D-alanine adduct indicates that following formation of PLP-D-alanine, the hydroxyl moiety of PLP forms a through-water hydrogen bond with the backbone carbonyl oxygen of Ser179 (26). Residue Ser179 in the *B. subtilis* Dat structure corresponds to Ser180 in *S. aureus*. If similar interactions mediate PLP binding with *S. aureus* Dat, an approximately 1 Å deviation of the position of the backbone oxygen caused by the S_180_F substitution may create steric hindrance between PLP and the peptide bond connecting residues 180 and 181. It is reasonable to suggest that this shift in backbone position affects the binding affinity of PLP to Dat when PLP is not covalently attached to Lys146. Additionally, the water mediated hydrogen bond between the residue 180 side chain and PLP is lost due to the S_180_F mutation.

There are additional minor conformational changes in the side chains of several other residues accommodating the added bulk of the Phe side chain in the S_180_F variant. The only prominent change to side chain structure is a shift of Lys157, which is pushed away from the active site due to steric hindrance with the Phe180 side chain. However, Lys157 does not appear to be important for substrate binding or catalysis. Other residues known to be important for Dat activity such as Tyr32, Arg99, and His101, which form the “carboxylate trap” in PLP-dependent enzymes catalyzing reactions on D-amino acids, position the carboxylate moiety of substrates to ensure correct binding orientation of substrate (26). The side chain conformations of the carboxylate trap residues are essentially unchanged in Dat-S_180_F when compared to wild-type Dat (Fig. 5D).

Taken together, the likely structural differences in the Dat-S_180_F variant are small but disproportionately relate to interactions between the Dat polypeptide and the PLP co-factor. This suggests that the S_180_F mutant has an altered affinity for the PLP-BCDA adduct, which would enable release of that adduct before the active site Lys146 can react with the BCDA side chain. While this would sacrifice a PLP molecule, Dat would remain able to bind another PLP molecule, form the covalent linkage between PLP and the Lys146 side chain, and inactivate additional BCDA. The underpinning chemistry of this possibility is somewhat similar to the known mechanism of BCDA inactivation in *M. tuberculosis* (24).

### Docking of β-chloro-D-alanine covalent adducts

The proposed mechanism of Dat inactivation by BCDA involves the initial formation of BCDA-PLP adduct followed by β-elimination of chloride via a 2-amino acrylate (2AA) external aldimine (2AA-PLP) intermediate (24, 27). Here molecular docking experiments were performed to assess possible affinity changes in the Dat-S_180_F variant using the BCDA-PLP adduct and the 2AA-PLP intermediate. Predictably, the BCDA-PLP docking results revealed similar binding modes for both proteins but with minor differences. BCDA-PLP docked in a similar orientation as pyridoxal D-alanine in complex with *B. subtilis* Dat (26). The docking results suggest that the carboxylate moiety of the BCDA-PLP adduct forms interactions with the carboxylate trap but lacks the bidentate interaction with Arg99 (Fig. 6B) (24, 25). The 2AA-PLP external aldimine also docked with a similar binding mode to that observed for the wild-type Dat crystal structure, but maintained a bidentate interaction between Arg99 and the carboxylate moiety of the 2AA-PLP also forming a hydrogen bond with the side chain of Tyr32 (26). Again, both residues are part of the carboxylate trap (RSCB PDB:3DAA). Additionally, Lys146 is positioned to afford a reaction with the 2AA-PLP, which would either simply reverse the initial reaction on PLP or, through multiple chemical steps, irreversibly inactivate the enzyme (24, 25). In contrast, the Dat-S_180_F docked models vary the position of the 2AA-PLP carboxylate moiety placing it near the carboxylate trap residues in some models but exhibiting a 180° rotation around the substrate amino acid φ bond that places the β-carbon in the region near the carboxylate trap, possibly preventing irreversible covalent modification of Lys146. Although unlikely due to the multiple interactions between the residues of the carboxylate trap and the α-carboxylate of the amino acid substrate, this dihedral rotation may become feasible as a result of β-elimination of chloride from the BCDA-PLP adduct and formation of the 2-AA external aldimine (Fig. 6C).

The structural and docking results suggest that differences in the PLP binding site caused by the S_180_F mutation likely modulate PLP-adduct affinity during the multi-step chemistry necessary for reaction with Lys146. In doing so, these structural differences may afford release of catalytic intermediates representing inactivated BCDA. Consistent with this the IC_50_ of Dat-S_180_F is lower than wild-type.

### Contribution of D-ala-D-ala pathway enzymes to BCDA resistance

Unlike the Dat-S_180_F allele in BCDA1, which as described above appears to retain enough catalytic activity to act on BCDA and chemically alter it, the *dat*::Em mutant is presumably incapable of inactivating BCDA and its MIC is only slightly reduced (Table 1). In contrast the BCDA MIC of *alr1* is reduced to <50 μg/ml (Table 1). These data suggest that either Alr1 activity can generate enough D-alanine to support growth in the presence of low BCDA concentrations even the absence of Dat and/or that BCDA may also be targeting the D-alanine ligase enzyme Ddl in the D-ala-D-ala pathway. To investigate this, the impact of exogenous D-alanine on the susceptibility of *alr1* and *dat*::Em to BCDA was measured. Consistent with its reduced MIC, growth of *alr1* was completely inhibited in CDM supplemented with BCDA 100 μg/ml, whereas the *dat*::Em mutant was capable of growing, albeit at reduced levels compared to JE2 (Fig. S2A). Exogenous D-alanine reversed the BCDA-mediated inhibition of growth of both *alr1* and *dat*::Em in a concentration dependent manner (Fig. S2B, C). Taken together these data reveal that in the absence of Alr1-mediated D-alanine synthesis, inhibition of Dat and Ddl by BCDA has a significant negative effect on growth. In contrast, in the absence of Dat activity, Alr1-mediated D-alanine synthesis enables growth at higher BCDA concentrations. The Dat-S_180_F mutation confers increased BCDA resistance not by altering D-alanine synthesis but via inactivation of the antibiotic.

## Discussion

Alanine metabolism plays a crucial role in cellular processes, including the regulation of cell wall biosynthesis, amino acid homeostasis, and energy production. Despite its importance, the enzymes involved in alanine metabolism remain relatively underexplored in *S. aureus*, particularly in the context of metabolic adaptation to antibiotic exposure and expression of resistance. Here we sought to advance our understanding of these pathways and their contribution to growth under physiologically relevant, nutrient-limited conditions to generate new insights into antibiotic resistance in MRSA.

Interconnected central metabolism and amino acid biosynthetic pathways are regulated in a manner dependent on the available carbon source(s). In the absence of its preferred carbon source, glucose, growth of *S. aureus* is supported by amino acids such as alanine and aspartate (28, 29). Our data showing the reversal of L-alanine auxotrophy by glucose illustrates the metabolic flexibility of *S. aureus* and the interplay between central metabolism and amino acid biosynthesis. The Alr1 and Dat enzymes, and not Ald1 or Ald2, are essential for growth in chemically defined medium (CDM) lacking alanine. Furthermore, suppressor mutants enabling growth in CDM lacking L-alanine were readily isolated but only if *alr1* and *dat* were intact. In CDM with glucose (CDMG), the *dat* mutant required either L- or D-alanine for growth, while the *alr1* mutant could only grow when exogenous L-alanine was available. This underscores the critical role of Alr1 not only for the synthesis of D-alanine, but also for the conversion of D-alanine to L-alanine under alanine-limited growth conditions. Interestingly, the Alr2 homologue cannot compensate for the absence of Alr1 and indeed no growth defect or change in DCS/β-lactam susceptibility was observed for the *alr2* mutant in CDM or CDMG.

Beyond Alr2, it remains unclear why alanine dehydrogenase 1 and 2 (Ald1 and Ald2), which are predicted to catalyze the conversion of pyruvate to L-alanine (Fig. 1), cannot compensate for the absence of Alr1 for the synthesis of L-alanine. Indeed our data revealed no growth defect for an *ald1*/*ald2* double mutant under any growth condition tested, which are largely consistent with a previous study in *S. aureus* which reported no detectable phenotype for *ald2* and an extended lag phase for *ald1* in CDM but no change in final cell density (16). These data raise questions about the functions of Ald1 and Ald2 in *S. aureus* compared to other microorganisms. Possibly Ald1 and Ald2 do not undertake this function in *S. aureus* or the enzymes are not expressed under these conditions. In *S. aureus ald1* forms an operon with *ilvA*, encoding threonine dehydratase, while *ald2* is located elsewhere in the genome. In *Paeniclostridium sabinae*, which also has two *ald* genes, *ald1*, but not *ald2*, facilitates alanine synthesis from ammonia and pyruvate (30). *Mycobacterium smegmatis* has a single *ald* gene implicated in alanine utilization and anaerobic growth (31). *Bacillus subtilis* also has a single alanine dehydrogenase gene, which is reported to support growth when alanine is the sole carbon source (32). Future characterization of the suppressor mutations that reverse alanine auxotrophy in CDM may provide insights on the pathways required for L-alanine biosynthesis under alanine-limited growth conditions, including the roles of *ald1* and *ald2*.

Our previous paper (14) and the data presented in this study underscore the importance of alanine transport and biosynthesis in antibiotic resistance. Consistent with earlier findings, mutation of *alr1* significantly increases susceptibility to oxacillin, DCS and BCDA in CDM. In contrast the *dat* mutation had no impact on oxacillin and DCS susceptibility, and only marginally reduced the BCDA MIC. Mutation of the alanine transporter *cycA* increased susceptibility to oxacillin and DCS, but not BCDA. These data indicate that DCS and BCDA, which are both alanine analogue antibiotics, have different mechanisms of action. One explanation for the observation that the *alr1* mutant was significantly more susceptible to BCDA than the *dat* mutant (MIC <50 μg/ml versus 200 μg/ml) is that BCDA targets the Dat enzyme. Because D-alanine biosynthesis in the *alr1* mutant is completely reliant on Dat activity, inhibition of Dat activity by BCDA significantly compromises D-alanine availability thereby exacerbating vulnerability to this antibiotic. We previously reported that alternative alanine transporter(s) can compensate for CycA in CDM (14), providing a possible explanation for why the D-alanine pathway is less perturbed in this mutant and why it is less vulnerable to BCDA-mediated inhibition of Dat. It is also notable that mutation of *dat* did not affect DCS susceptibility, indicating that Dat does not significantly contribute to production of D-alanine when Alr1 is active. This conclusion supports a previous report that Dat is more active in the conversion of D-alanine to pyruvate, rather than vice versa (19). Finally, aligned to their lack of impact on growth, *alr2*, *ald1* and *ald2* mutations did not affect susceptibility to OX, DCS and BCDA, further excluding these enzymes from a significant role in the D-alanine pathway and peptidoglycan biosynthesis.

Supporting the possibility that Dat is a target for BCDA, a novel C_539_T mutation in the *dat* gene was identified as the only genetic change in a BCDA resistant mutant of JE2. Swapping the *dat*_C539T_ allele with *dat*::Em reversed the increase in BCDA resistance, while multi-copy expression of the *pepV*-*dat*_C539T_ operon in wild-type MRSA significantly increased BCDA resistance. In *Bacillus sphaericus* BCDA binds to and inactivates D-amino acid transaminase (D-AAT) (33). D-AAT has also been shown to bind to β-cyano-D-alanine, another D-alanine derivative, leading to inactivation of the enzyme (34). These studies suggest that BCDA may act as a competitive substrate and/or inhibitor of Dat in *S. aureus*. Dat functions as a bidirectional enzyme, catalyzing the reversible conversion of D-alanine to pyruvate. *In vitro* enzyme assays revealed that production of pyruvate by Dat-S_180_F was reduced to 43% of wild-type levels. Consistent with this, a R_179_G substitution in the active site of the *Haliscomenobacter hydrossis* DAAT also decreased enzyme function (35). Furthermore the strict D-alanine auxotrophy of both the *alr1*/*dat*::Em and *alr1*/*dat*_C539T_ double mutants support the conclusion that the Dat-S_180_F enzyme cannot compensate for the loss of Alr1 to meet the requirement for D-alanine synthesis.

While the acquisition of resistance to BCDA through a mutation in *dat* seems logical, the more intriguing question is why or how this resistance is accompanied by altered Dat transamination activity? Given the hypothesis that BCDA targets Dat, the identification of a mutation in the *dat* gene leading to BCDA resistance was not unexpected. However the data showing that this mutation reduced the normal ability of the enzyme to catalyze the conversion of D-alanine to pyruvate and that the IC_50_ of Dat-S_180_F for BCDA was significantly lower than wild-type Dat raises the intriguing question of how the S_180_F substitution in Dat leads to BCDA resistance. Dat is a pyridoxal phosphate (PLP)-dependent enzyme and we determined the X-ray crystal structure of Dat in complex with PLP to gain insight into its catalytic mechanism and cofactor interactions. Structural comparisons with *Bacillus sp.* YM-1 Dat showed that the active site residues, including those interacting with PLP, are conserved, indicating the functional importance of this region in catalysis. Further, the structural comparison between wild-type Dat and the S_180_F variant highlighted a subtle but functionally important shift in the active site loop, likely affecting the affinity of PLP binding. This finding aligns with a previous study on D-AAT, in which a Y_31_Q substitution was associated with significantly reduced enzyme activity and enhanced susceptibility to BCDA (36). A E_177_K mutation in D-AAT also significantly impacted coenzyme anchoring and catalytic efficiency (37) further emphasizing the significance of coenzyme binding site residues in enzyme stability and activity. Furthermore, these D-AAT mutations can alter the enzyme’s stereochemical fidelity, increasing its ability to convert L-alanine to D-alanine. This has not been examined in our study and may be important.

Molecular docking analysis identified a possible mechanism through which the S_180_F substitution in Dat may change how the enzyme interacts with the BCDA-PLP adduct, leading to BCDA resistance. Thus, while the Dat_ Dat-S_180_F variant exhibits a reduced capacity to catalyze the reversible conversion of D-alanine to pyruvate, the IC_50_ of the mutant enzyme for BCDA is decreased, which is consistent with the molecular docking analysis suggesting that Dat-S_180_F can bind BCDA, form the BCDA-PLP adduct, and release an inactivated form of BCDA, such as 2AA-PLP. If 2AA-PLP is released by Dat-S_180_F, this product would likely react with water to form pyruvate-PLP. Further validation of this predicted mechanism will be needed. Given the crucial role of PLP-dependent enzymes in bacterial physiology and antibiotic resistance, it would be interesting to investigate if similar resistance mechanisms exist in other bacterial species and if targeting the PLP interactions using modified BCDA could offer new strategies for overcoming BCDA resistance in *S. aureus*.

Taken together our data reveal essential roles for Alr1 and Dat in alanine metabolism, growth and antibiotic resistance in MRSA. The ability of exogenous D-alanine to reverse growth inhibition of the *alr1* and *dat*::Em mutants by BCDA suggests that BCDA may competitively inhibit the Ddl D-alanine ligase enzyme as well as Dat. In contrast DCS targets Alr1 and Ddl revealing that these antibiotics together inhibit the three major enzymes in the D-ala-D-ala pathway. Consistent with this we report synergy between DCS and BCDA against MRSA and propose that their combined use may limit the emergence of resistance and have significant therapeutic potential, particularly if used in conjunction with β-lactams.

## Experimental procedures

### Bacterial strains, reagents and growth conditions

The bacterial strains and plasmids used in this study are listed in Table S1. *Staphylococcus aureus* strains were grown in Tryptic Soy broth (TSB), Tryptic Soy agar (TSA), Mueller Hinton agar (MHA), chemically defined media (CDM) and CDM supplemented with glucose (5 g/L) (CDMG) (16). CDM and CDMG were further manipulated to remove L-alanine (CDM/CDMG - L-ala), to replace L-alanine with 5mM D-alanine (CDM/CDMG - L-ala + D-ala) or to add 5mM D-alanine (CDM/CDMG + L-ala + D-ala). *Escherichia coli* strains IMO8B, XL-1 Blue, and BL21 (DE3) were cultured in Luria Bertani broth (LB) or Agar (LBA). *S. aureus* JE2 mutants from the Nebraska Transposon Mutant library (NTML) used in this study were NE810 *cycA*::Em, NE16 *ald1*::Em, NE198 *ald2*::Em, NE1713 *alr1*::Em, NE799 *alr2*::Em and NE1305 *dat*::Em. *E. coli*-*S. aureus* shuttle plasmid pLI50 (38) was used for complementation analysis and plasmid pET28b was used for overexpression and purification of recombinant proteins. Growth media were supplemented with appropriate antibiotics, including β-chloro-D-alanine (BCDA, 500 µg/ml), erythromycin (Em, 10 µg/ml), chloramphenicol (Cm, 10 µg/ml), spectinomycin (Spc, 1000 µg/ml), kanamycin (Km, 50 µg/ml), and ampicillin (Amp, 100 µg/ml), oxacillin (OX) and D-cycloserine (DCS) as required.

### Mutant strain construction

The *ald1*::Em, *ald2*::Em, and *dat*::Em strains obtained from NTML were used to construct *ald1*::Em/*dat*::Spc and *ald2*::Em/*dat*::Spc double mutants, as well as the *ald1*::Em/*ald2*::Km/*dat*::Spc triple mutant. To generate double mutants, the Em antibiotic cassette in the NE1305 *dat*::Em transposon mutant was first replaced with Spc via allelic exchange using the plasmid pSPC, which harbours a temperature-sensitive origin as described previously (39). This was followed by transduction using a lysate prepared from the *ald1*::Em *and ald2*::Em alleles were then transduced into the *dat*::Spc mutant using phage 80α. To construct the *ald1*::Em/*ald2*::Km/*dat*::Spc triple mutant, the pKAN plasmid was used to generate the *ald2*::Km mutant. The *ald2*::Km was transduced into *dat*::Spc, to generate the *ald2*::Km/*dat*::Spc double mutant, into which the *ald1*::Em allele was transduced.

The *dat*::Spc was transduced with a lysate prepared from the *alr1*::Em mutant to generate an *alr1*/*dat* double mutant. Lysates from the *alr1*::Em and *alr2*::Em mutants were also transduced into BCDA1 to generate *alr1*/*dat_C539T_* and *alr2*/*dat_C539T_* mutants, respectively. All double and triple mutants were confirmed by PCR amplification of the target loci using primers listed in Table S2.

### Growth assays in CDM and CDMG

The growth of *S. aureus* strains was measured using a Tecan Sunrise microplate reader, with data recorded and analyzed via Magellan software. In brief, overnight cultures of each strain grown in TSB supplemented with appropriate antibiotics were harvested by centrifugation at 10,000 x g for 5 min, washed twice in phosphate-buffered saline (PBS), and resuspended in PBS to an OD_600_ = 1. A 10 µl of this suspension was inoculated into 190 µl of the respective medium in 96-well hydrophobic polystyrene plates, resulting in a final volume of 200 µl with a starting OD_600_ = 0.05. Plates were then incubated with shaking at 35-37 □ for 18-24 hours in a pre-heated Tecan microplate reader, with the OD_600_ recorded every 15-minutes. Growth assays were performed using at least three independent biological replicates, and the resulting data was plotted using GraphPad Prism software.

### Generation of a BCDA-resistant mutant

To generate a BCDA-resistant mutant, a TECAN growth assay was performed using wild-type JE2. Briefly, overnight cultures were washed and resuspended in PBS at an OD_600_ = 1, from which 10 µl was used to inoculate a 96-well plate containing 190 µl of CDM supplemented with BCDA at concentrations of 1, 50 and 500 µg/ml. The MIC of BCDA for JE2 is 300 µg/ml. Growth was monitored for 24 hours as described above. Growth was observed at 1 and 50 µg/ml BCDA, whereas 500 µg/ml BCDA inhibited growth in most wells. However, one of the wells inoculated with JE2 at 500 µg/ml showed visible growth after 24 h, from which single colonies were isolated on TSA. The BCDA resistant phenotype of one isolated colony was confirmed by TECAN growth assays, and designated BCDA1. Following whole genome sequencing analysis (described below), the *dat*_C539T_ allele of BCDA1 was replaced with *dat*::Em from NE1305 by phage 80α transduction as described above, and verified using FP_*dat* and RP_*dat* primers (Table S2).

### Genomic DNA (gDNA) extraction and whole genome sequencing (WGS)

Cell pellets from 3 ml of overnight *S. aureus* cultures were pre-treated with lysostaphin (10 µg/ml; Ambi Products LLC) at 37 □ for 30 minutes prior to gDNA extraction using the Wizard Genomic DNA Purification Kit (Promega) (40), and sequencing was conducted by MicrobesNG using Illumina sequencing platform with 2×250 bp paired end reads. Genome sequence analysis was performed as described previously (41) using CLC Genomics Workbench software (Qiagen, Version 22.0.1). The *S. aureus* JE2 genome sequence was used as a reference, with a contig produced by mapping Illumina reads onto the closely related USA300_FPR3757 genome sequence (RefSeq accession number NC_007793.1). The Illumina short-read sequence of the BCDA1 strain was then compared to the assembled JE2 sequence to identify single nucleotide polymorphisms (SNPs), insertions, or deletions.

### Antibiotic minimum inhibitory concentration (MIC) and fractional inhibitory concentration (FIC) measurements

MIC measurements by broth microdilutions were performed in accordance with CLSI methods for dilution susceptibility testing of staphylococci (42) with modifications. Briefly, strains were first grown at 37°C on MHA for 24 h and 5 - 10 colonies were resuspended in 0.85% saline before being adjusted to 0.5 McFarland standard (*A*_600_ = 0.1). The cell suspension was then diluted 1:20 in PBS and 10 ml used to inoculate 100 ml culture media (MHB or CDM) containing serially diluted antibiotics in 96-well plates.

To investigate synergy between DCS and BCDA the FIC index (ΣFIC) for MRSA strain JE2 was calculated. The ΣFIC = FIC A + FIC B, where FIC A is the MIC of DCS in combination with BCDA / the MIC of DCS alone, and FIC B is the MIC of BCDA in combination with DCS / the MIC of BCDA alone. Antibiotic combinations are considered synergistic when the ΣFIC is ≤0.5, indifferent when the ΣFIC is >0.5 to <2 and antagonistic when the ΣFIC is >2.

### Complementation of the BCDA1 mutant

To complement the BCDA1 mutant, the *pepV*-*dat* operon and upstream regulatory sequences from JE2 and BCDA1 were amplified and cloned into plasmid pLI50 using *EcoRI* and *SalI* restriction sites, generating recombinant plasmids p*pepV*-*dat*, and p*pepV*-*dat*_C539T_. For control purposes the *pepV* gene alone from JE2 was also amplified and cloned into pLI50 (38) to generate p*pepV*. These plasmids were first transformed into electrocompetent *E. coli* IMO8B cells, verified by Sanger sequencing (Eurofins Genomics) and subsequently transformed into JE2 and BCDA1 by electroporation.

### Expression and purification of recombinant *S. aureus* Dat in *E. coli*

The *dat* and *dat*_C539T_ genes were amplified from gDNA isolated from JE2 and BCDA1, respectively, using FP_Dat and RP_Dat primers (Table S2). The PCR products were digested with *EcoR*I and *Sal*I and ligated into the pET28b plasmid. The resulting recombinant plasmids, pET28b_Dat and pET28b_Dat_C539T_, were verified by Sanger sequencing (Eurofins Genomics) and transformed into *E. coli* BL21 (DE3). The BL21 cells containing a pET plasmid harboring the wild-type Dat enzyme were grown at 37°C in Luria broth media until OD_600_ reached 0.6. 1 mM IPTG was added to induce protein production at 16 °C for 20 hours. The cells were harvested through centrifugation at 3756 x *g*. They were resuspended in 20 mM Hepes pH 7.5, 500 mM NaCl, 5 mM β-mercaptoethanol, and 5 mM Imidazole buffer. Cells were lysed and DNA degraded by adding lysozyme (Hampton Research) and DNase I (Roche) and incubated on ice for 30 minutes. Cells were further lysed by sonication (Sonicator 3000, Misonix) and crude cell lysate was separated by centrifugation at 16,000 × *g* (Fixed angle rotor, 5810-R centrifuge, Eppendorf). The supernatant was applied to a 5 mL HisTrap TALON crude cobalt column (Cytiva). After washing with running buffer (20 mM Hepes pH 7.5, 500 mM NaCl, 5 mM β-mercaptoethanol, and 5 mM imidazole) to remove unbound protein, the recombinant Dat was eluted using a similar buffer as running buffer with 150 mM imidazole. The protein was further purified by size exclusion chromatography using 20 mM Hepes pH 7.5, 150 mM NaCl and 0.5 mM Tris (2-carboxyethyl) phosphine (TCEP) buffer as the mobile phase. Fractions containing purified Dat were pool and 1 mM PLP was added and excess PLP was removed by ultrafiltration with a buffer composed of 20 mM Hepes pH 7.5, 150 mM NaCl and 0.5 mM TCEP buffer.

### BCDA inhibition of Dat enzyme kinetics

Wild-type Dat and the Dat-S_180_F variant were purified as indicated above. To assess Dat enzymatic production of pyruvate from D-alanine, Dat enzymatic activity was coupled to that of Lactate Dehydrogenase (LDH). The reactions were monitored by continuously measuring absorbance at 340 nm using the Synergy H4 Hybrid Reader from BioTek. The assay component final concentrations were 50 nM DAT (WT or S_180_F), 100 mM Hepes pH 7.5, 10 mM α-ketoglutarate, 400 µM NADH, 25 mM D-alanine, 1.2 U LDH and 10 µM PLP. The inhibition of Dat by BCDA was determined using various concentrations of BCDA and the respective IC_50_ values of BCDA against WT and S_180_F Dat were fit using GraphPad Prism software.

### Crystallization, X-ray diffraction and data processing

The vapor diffusion hanging drop method was used for crystallization experiments. The Dat protein was in 20 mM Hepes pH 7.5, 150 mM NaCl and 0.5 mM TCEP buffer at 7 mg/mL concentration. The protein and well solution were mixed at a 1:1 volumetric ratio and equilibrated against the well solution containing 0.1 M BIS-TRIS pH 6.5, 45% v/v Polypropylene glycol P 400. The harvested crystals were flash cooled in liquid nitrogen. The X-ray diffraction experiments were performed at the University of Nebraska Medical Center, Structural Biology Core Facility.

The diffraction data were processed in CCP4 using DIALS. The crystal structure of D-amino acid aminotransferase/PLP of *Bacillus* sp. YM-1 (RCSB PDB code: 1DAA) was used for molecular replacement in PHENIX (43). The PLP was modeled using eLBOW and the covalent adduct was made using PHENIX (44). PHENIX was used for the structure refinement and iterated with manual model building using Coo (45, 46). The structure was validated using Molprobity (47).

### Molecular docking experiments

Using a biological dimer derived from the *S. aureus* Dat X-ray crystal structure both wild-type Dat and the Dat-S_180_F variant were modeled in Glide (Schrödinger) using energy minimization. Schrödinger LigPrep was used to prepare the covalent adducts formed by the reaction of PLP and BCDA. Molecular docking experiments employed the PLP-BCDA adducts following receptor grid preparation for both the wild-type and S_180_F variant. The results were exported to PyMOL and visually evaluated.

## Supporting information

wwPDB full validation report for Dat X-ray crystal structue

## ACKNOWLEDGEMENTS

This work was supported by Research Ireland grant 19/FFP/6441 to J.P.O’G, Research Ireland Pathway Award 22/PATH-S/10804 to M.S.Z. and NIH/NIAID R01AI125588 to V.C.T. The funders had no role in the study design, data collection, interpretation, writing of the manuscript, or the decision to publish.

Conceptualization: R.R., Y.J., M.S.Z., D.R.R. and J.P.O’G. Methodology: R.R., S.P., Y.J., and S.P. Formal analysis: R.R., J.Y., M.S.Z, V.C.T., D.R.R. and J.P.O’G. Writing-original draft: R.R., Y.J., D.R.R. and J.P.O’G. Writing-review, editing and approval: all authors. Funding acquisition: J.P.O’G, M.S.Z. and V.C.T. Project administration: J.P.O’G.

## Data Availability

The Full Worldwide Protein Data Bank (wwPDB) X-ray Structure Validation Report for *Staphylococcus aureus* D-alanine aminotransferase in complex with pyridoxal 5’-phosphate is available under accession ID: 9PXS / pdb_00009pxs.

---

**Fig. S1. DCS and BCDA are synergistic against MRSA. A.** Disk diffusion assays with DCS (30 μg) and BCDA (1000 μg) against JE2 grown on Mueller-Hinton agar for 24 h at 37°C. **B.** Checkerboard titration assays conducted using DCS and BCDA with JE2 grown for 24 h at 37 °C in Mueller-Hinton broth in 96-well plates. The data shown are the OD_600_ values for each well. The experiments were repeated at least three times and the data from a representative 96-well plate is shown. Red shaded boxes indicated wells in which growth was measured.

**Fig. S2. Exogenous D-alanine restores growth of *alr1* and *dat*::Em mutants inhibited by BCDA. A.** Comparison of JE2, BCDA1, NE1305 (*dat*::Em) and *alr1* growth in CDM supplemented with BCDA 200 μg/ml. **B.** Comparison of *alr1* growth in CDM supplemented with BCDA 100 μg/ml alone or with exogenous D-alanine concentrations from 0.005 to 5 mM. **C.** Comparison of *dat*::Em growth in CDM supplemented with BCDA 500 μg/ml alone or with exogenous D-alanine concentrations from 0.005 to 5 mM. The data presented are the average of at least 3 biological replicates and standard deviations are shown.

