## Supplementary material for "A D-alanine aminotransferase S_180_F substitution confers resistance to β-chloro-D-alanine in *Staphylococcus aureus* via antibiotic inactivation": Fig. S1

A

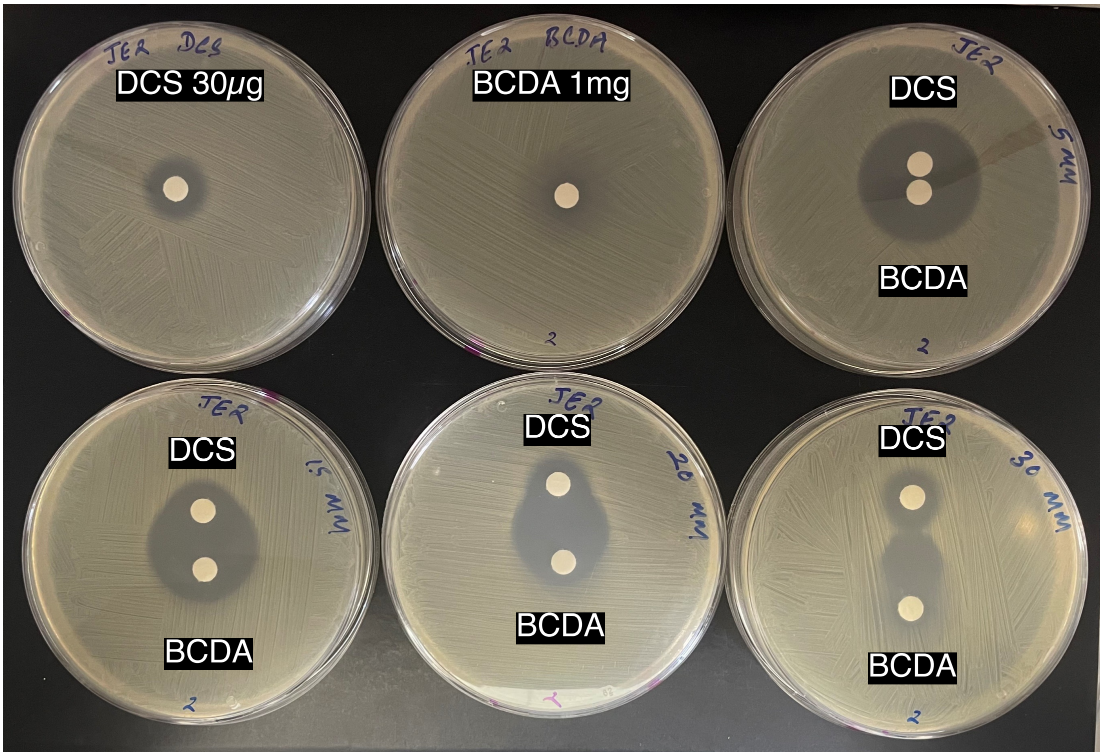


B

| DCS (μg/ml) | | | | | | | | | | | | |
|---|---|---|---|---|---|---|---|---|---|---|---|---|
| 128 | 0.076 | 0.075 | 0.073 | 0.072 | 0.074 | 0.075 | 0.073 | 0.074 | 0.076 | 0.078 | 0.077 | 0.086 |
| 64 | 0.075 | 0.076 | 0.075 | 0.073 | 0.074 | 0.074 | 0.073 | 0.074 | 0.076 | 0.077 | 0.076 | 0.08 |
| 32 | 0.073 | 0.075 | 0.083 | 0.074 | 0.076 | 0.078 | 0.074 | 0.075 | 0.078 | 0.078 | 0.077 | 0.082 |
| 16 | 0.339 | 0.233 | 0.171 | 0.157 | 0.083 | 0.075 | 0.074 | 0.074 | 0.076 | 0.076 | 0.077 | 0.081 |
| 8 | 0.508 | 0.442 | 0.407 | 0.399 | 0.251 | 0.075 | 0.095 | 0.074 | 0.077 | 0.076 | 0.076 | 0.079 |
| 4 | 0.502 | 0.425 | 0.423 | 0.457 | 0.319 | 0.312 | 0.088 | 0.076 | 0.079 | 0.079 | 0.078 | 0.085 |
| 2 | 0.651 | 0.455 | 0.451 | 0.429 | 0.422 | 0.373 | 0.218 | 0.087 | 0.077 | 0.079 | 0.079 | 0.083 |
| 0 | 0.637 | 0.441 | 0.428 | 0.416 | 0.414 | 0.438 | 0.384 | 0.39 | 0.298 | 0.203 | 0.182 | 0.083 |
| JE2 | 0 | 2 | 4 | 8 | 16 | 32 | 64 | 128 | 256 | 512 | 1024 | 2048 |

BCDA (μg/ml)

**Fig. S1. DCS and BCDA are synergistic against MRSA. A.** Disk diffusion assays with DCS (30 μg/ml) and BCDA (1000 μg/ml) against JE2 grown on Mueller-Hinton agar for 24 h at 37°C. **B.** Checkerboard titration assays conducted using DCS and BCDA with JE2 grown for 24 h at 37°C in Mueller-Hinton broth in 96-well plates. The data shown are the $OD_{600}$ values for each well. The experiments were repeated at least three times and the data from a representative 96-well plate is shown. Red shaded boxes indicated wells in which significant growth was measured.
