## Supplementary material for "A D-alanine aminotransferase S_180_F substitution confers resistance to β-chloro-D-alanine in *Staphylococcus aureus* via antibiotic inactivation": Fig. S2

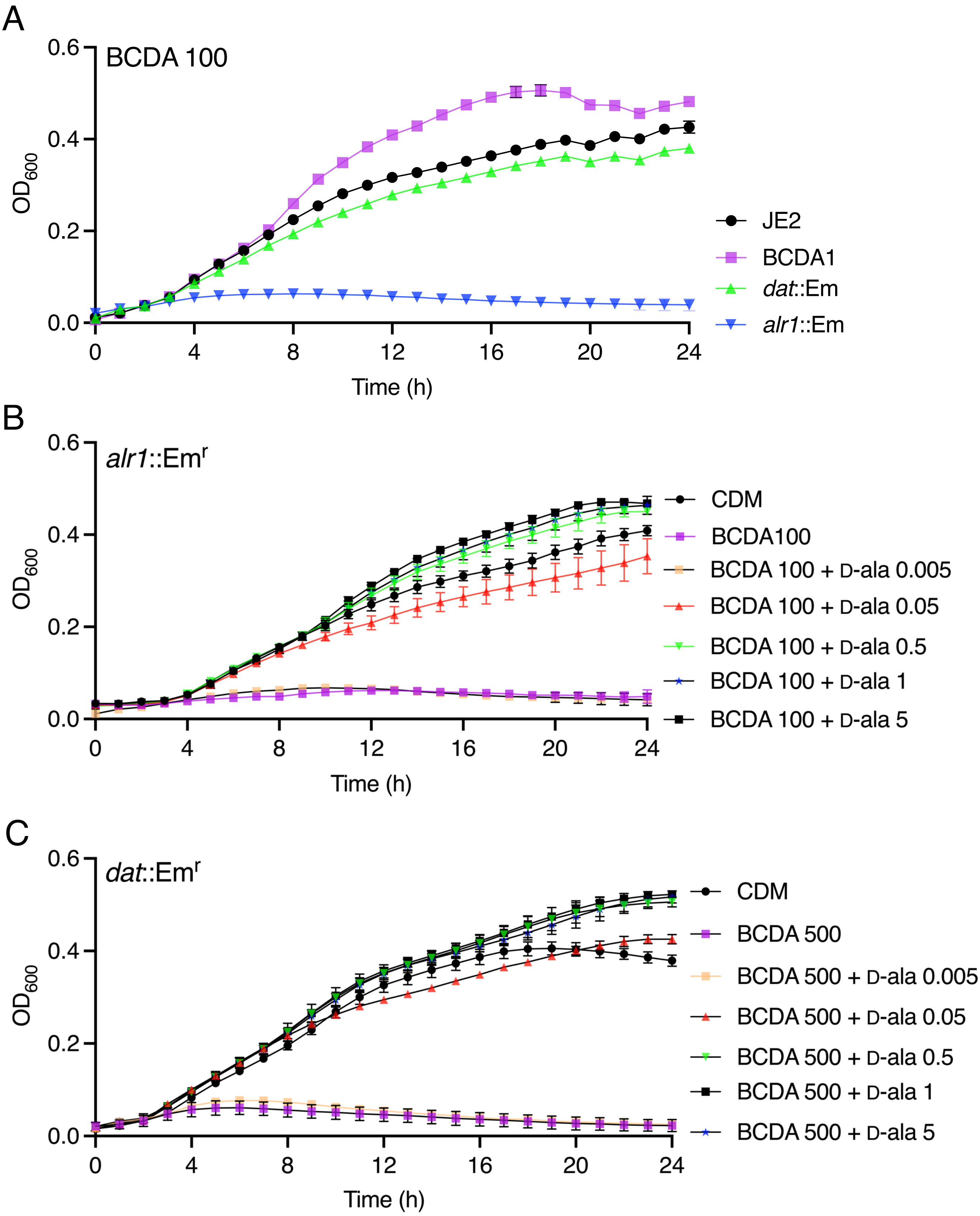


**Fig. S2. Exogenous D-alanine restores growth of *alr1* and *dat*::Em$^r$ mutants inhibited by BCDA. A.** Comparison of JE2, BCDA1, NE1305 (*dat*::Em) and *alr1* growth in CDM supplemented with BCDA 100 µg/ml. **B.** Comparison of *alr1* growth in CDM supplemented with BCDA 100 µg/ml alone or with exogenous D-alanine concentrations from 0.005 to 5 mM. **C.** Comparison of *dat*::Em$^r$ growth in CDM supplemented with BCDA 100 µg/ml alone or with exogenous D-alanine concentrations from 0.005 to 5 mM. The data presented are the average of at least 3 biological replicates and standard deviations are shown.
