## Supplementary material for "A D-alanine aminotransferase S_180_F substitution confers resistance to β-chloro-D-alanine in *Staphylococcus aureus* via antibiotic inactivation": Table S2

**Table S2: Primers used in this study**

| Primer name | Sequence (5’-3’) | Purpose | Restriction site |
| --- | --- | --- | --- |
| FP_*cycA* | ACAGAATAGCCACAAATAGCACC | *cycA* mutant verification | - |
| RP_*cycA* | GAACTTAATGTCCCAAGCCCT |  | - |
| FP_*ald1* | GAGGTGTTCGACGAATAAATGG | *ald1* mutant verification | - |
| RP_*ald1* | TTCACAAATGAAAGAGGAGTGTTGC |  | *-* |
| FP_*ald2* | CGCTGTAATCCATTCCCTTT | *ald2* mutant verification | - |
| RP_*ald2* | TGAACCGTCAACTGCGATTA |  | - |
| FP_*dat* | GGATCCGACAAGGGTGTAGCATTTGGC | Verification of BCDA1 strain | *Bam*HI |
| RP_*dat* | GAATTCCCTCAACCAATGCCTACATTACACG |  | *Eco*RI |
| FP_Dat | CGGAATTCGGAAAAAATTTTTTTAAATGGTGAGTTTGTAAGTCC | Recombinant Dat over-expression and purification | *Eco*RI |
| RP_Dat | ACGCGTCGACTTAAATACTGTGTGACTCTATATACTTTTCAAATCCTTC |  | *Sal*I |
| FP_*pepV-dat* | ACGCGTCGACGCCAACTTGCATTGTCTGTAG | Complementation of *dat* mutations | *Sal*I |
| RP_*pepV* | CGGAATTCCACTTGGACTTACAAACTCACC |  | *Eco*RI |
| RP_*pepV-dat* | CGGAATTCCCTCAACCAATGCCTACATTACACG |  | *Eco*RI |
| FP_*alr1* | CGGAAAAGCTTCGTTCGCTAGG | *alr1* mutant verification | **-** |
| RP_*alr1* | CAACGACGAACTTGGCAAACC |  | **-** |
| FP_*alr2* | CGCCATTCTTATTTGGGGAAGA | *alr2* mutant verification | **-** |
| RP_*alr2* | TGGTAACGGCGCATCTGTTC |  | **-** |
| FP_T7-promoter | TAATACGACTCACTATAGGG | pET28b recombinant plasmids | **-** |
| RP_T7-terminator | GCTAGTTATTGCTCAGCGG |  | **-** |
