## Supplementary material for "A D-alanine aminotransferase S_180_F substitution confers resistance to β-chloro-D-alanine in *Staphylococcus aureus* via antibiotic inactivation": Table S1

**Table S1:** Strains and plasmids used in this study

| Strains or plasmids | Description | Source |
| --- | --- | --- |
| Strains |  |  |
| *S. aureus* JE2 | CA-MRSA, USA300, cured of plasmids. Parent strain of Nebraska Transposon Mutant library (NTML). | (1) |
| NE810 | NTML mutant. *cycA*::Em^r^ | (1) |
| NE1713 | NTML mutant. *alr1*::Em^r^ | (1) |
| NE799 | NTML mutant. *alr2*::Em^r^ | (1) |
| NE1136 | NTML mutant. *ald1*::Em^r^. | (1) |
| NE198 | NTML mutant. *ald2*::Em^r^. | (1) |
| NE1305 | NTML mutant. *dat*::Em^r^ | (1) |
| NE1898 | NTML mutant. *pepV*::Em^r^ | (1) |
| BCDA1 | BCDA resistant derivative of JE2. C-to-T mutation at nucleotide 539 of the *dat* gene (SAUSA300_1696) resulting in a predicted S_180_F amino acid substitution in Dat. | This study |
| BCDA1 *dat*::Em^r^ | *dat*::Em^r^ transduced into BCDA1 strain | This study |
| JE2 pLI50 | Plasmid pLI50 transformed into JE2 | This study |
| JE2 p*pepV* | Plasmid p*pepV* transformed into JE2 | This study |
| JE2 p*pepV*-*dat* | Plasmid p*pepV*-*dat* transformed into JE2 | This study |
| JE2 p*pepV*-*dat*_C539T_ | Plasmid p*pepV*-*dat*_C539T_ transformed into JE2 | This study |
| BCDA1 p*pepV*-*dat* | Plasmid p*pepV*-*dat* transformed into JE2 | This study |
| *alr1/dat*_C539T_ | *alr1*::Em allele transduced into BCDA1 strain | This study |
| *alr2/dat*_C539T_ | *alr2*::Em allele transduced into BCDA1 strain | This study |
| *dat*::Spc | *dat*::Em^r^ allele in NE1305 swapped for *dat*::Spc | This study |
| *ald2*::Km | *ald2*::Em allele in NE198 swapped for *ald2*::Km | This study |
| *alr1*::Em/*dat*::Spc | *alr1*::Em allele transduced into *dat*::Spc strain | This study |
| *dat*::Spc/*ald1*::Em | *ald1*::Em allele transduced into *dat*::Spc strain | This study |
| *dat*::Spc/*ald2*::Em | *ald2*::Em allele transduced into *dat*::Spc strain | This study |
| *dat*::Spc/*ald2*::Km | *ald2*::Km allele transduced into *dat*::Spc strain | This study |
| *ald1*::Em/*ald2*::Km/*dat*::Spc | *ald1*::Em allele transduced into *dat*::Spc/*ald2*::Km strain | This study |
| *E. coli* XL-1 Blue | General plasmid maintenance strain | Agilent Technologies |
| *E. coli* BL21 (DE3) | Strain for protein over-expression and purification | Invitrogen |
| *E. coli* BL21 (DE3) pET28b | Plasmid pET28b transformed into *E. coli* BL21 (DE3) | This study |
| *E. coli* BL21 (DE3) pET28b_Dat | Plasmid pET28b_Dat transformed into *E. coli* BL21 (DE3) | This study |
| *E. coli* BL21 (DE3) pET28b_*dat*_C539T_ | Plasmid pET28b_*dat*_C539T_ transformed into *E. coli* BL21 (DE3) | This study |
| *E. coli* IM08B | *E. coli* DC10B with the staphylococcal (CC8-2)-type methylation system integrated between *atpI* and *gidB* and the (CC8-1)-type methylation system integrated between *essQ* and *cspB.* | (2) |
| Plasmids |  |  |
| pLI50 | *E. coli*-*S. aureus* shuttle vector | (3) |
| pET28b | Vector for protein over-expression and purification in *E. coli* BL21 (DE3) | Novagen |
